# Quantifying Uncertainty in Brain Age Predictions via Conformal Prediction

**DOI:** 10.64898/2026.09.04.749442

**Authors:** Ujjwal Samanta, Golia Shafiei, Helmet Karim, Theodore D. Satterthwaite, Andrew Chen

**Affiliations:** Department of Public Health Sciences, Medical University of South Carolina, Charleston, SC, USA; Penn Lifespan Informatics and Neuroimaging Center (PennLINC), Department of Psychiatry, Perelman School of Medicine, University of Pennsylvania, Philadelphia, PA, USA; Department of Psychiatry, University of Pittsburgh, Pittsburgh, PA, USA; Penn-CHOP Lifespan Brain Institute, Perelman School of Medicine, Children’s Hospital of Philadelphia Research Institute, Philadelphia, PA, USA; Department of Bioengineering, University of Pittsburgh, Pittsburgh, PA, USA

**Keywords:** Brain age, Conformal prediction, Uncertainty, MRI, Machine learning, Psychopathology, Youth, Prediction intervals, Neurodevelopment, Calibration

## Abstract

Youth can exhibit signs of delayed or accelerated neurodevelopment, measurable via brain magnetic resonance imaging (MRI). Brain age prediction seeks to estimate brain age at an individual level using machine learning models fitted on healthy individuals. The brain age gap (BAG)–the difference between brain age and chronological age–has been studied as a potential biomarker. However, BAG suffers from known limitations including dependence on chronological age, regression to the mean, and challenges in interpretability. As an alternative framework, we introduce brain age intervals (BAIs) as a normative interval on an individual’s chronological age, based on MRI measures. BAIs can be derived using a recent statistical framework called conformal prediction that yields prediction intervals with guaranteed coverage. We train several brain age prediction models on structural and functional MRI scans from the Reproducible Brain Charts dataset (aged 6-22) and estimated BAIs. The BAIs demonstrated stable empirical coverage close to the nominal 90% level (median coverage=91%, width=7.36 years, root mean square error (RMSE)=2.20 years across 100 repeated train–test splits). Association between interval coverage and clinical measures (p-factor, parental education) were weak, with only a modest directional signal for parental education. Together, these findings demonstrate the feasibility of uncertainty-aware brain age modeling in youth populations while highlighting the need for larger and more diverse samples to detect meaningful clinical and environmental associations.

## Introduction

Neurodevelopment is characterized by widespread changes in the brain, including cortical thinning, gray-matter reduction, and development of functional networks (1, 2). Large neuroimaging studies have revealed population-level trajectories of typical neurodevelopment. However, individuals are known to exhibit different rates of age-related changes in both brain structure and function (3). To identify deviations in neurodevelopment related to eventual psychopathology, it is essential to understand individual differences in neurodevelopment (4).

Brain age prediction models utilize neuroimaging to predict an individual’s age from structural and functional brain features derived from magnetic resonance imaging (MRI) (5). The difference between predicted brain age and chronological age—often referred to as the brain age gap (BAG)—has emerged as a promising biomarker of brain health and has been linked to neurological disorders, cognitive decline, and mortality risk (6-8). Brain age prediction approaches have also been applied to youth. Initial research showed that structural neuroanatomy predicts biological maturity independent of chronological age (9). Functional connectivity patterns also shift systematically over development, allowing machine learning classifiers to distinguish children from adults using resting-state connectivity alone with approximately 91-93% accuracy across independent datasets (10).

Even with the considerable advancements made in brain age prediction, various methodological issues persist. Existing models lack interpretability and instead concentrate mainly on prediction accuracy (5, 7). Paradoxically, as brain age model’s accuracy improves, its predictions approach chronological age, diminishing the biological signal that the brain age gap is meant to capture. Moreover, existing brain age models yield predicted age without quantifying uncertainty linked to those predictions (11, 12), leaving it unclear whether observed brain age difference reflect meaningful biological variation or merely model variability (7, 13). BAG in particular has major limitations, including correlation with chronological age and bias from regression towards the mean, which complicate its interpretation as an individual biomarker (7, 13). Recent work further demonstrates that summarizing brain age with single scalar loses substantial information and brain age derived models trained on chronological age may underperform models trained directly for specific clinical outcomes (21). With the growing interest in brain age for mapping individual deviations from typical neurodevelopment, there is a need for methods that capture uncertainty in brain age predictions and yield interpretable metrics (7, 12-14).

Current brain age prediction models depend on point estimates of estimated age without accounting for prediction uncertainty (5, 7, 12). Conformal prediction provides a model-agnostic, distribution-free approach for creating prediction intervals that ensure valid coverage (11, 15). By producing prediction intervals for brain age estimates, conformal prediction can inform whether an individual’s chronological age is within expected variability based on their MRI measures or indicate significant deviations from typical neurodevelopment. Incorporating conformal prediction may therefore improve the interpretability and reliability of brain age.

In this work, we apply conformal prediction to brain age prediction models to build individualized prediction intervals, which we call brain age intervals (BAIs). We examine whether falling outside these intervals relates to psychopathology and parental educational background in a large sample of youth from the Reproducible Brain Charts (RBC) (2). We compare several prediction models and conformal prediction approaches to identify intervals that achieve valid coverage with minimal width. As described below, our findings reflect both the potential of uncertainty-aware brain age frameworks and the challenges that remain in linking prediction uncertainty to meaningful individual differences

## Methods

### Dataset description

The RBC dataset (2) combined standardized neuroimaging and psychiatric profiling from five extensive developmental cohorts – the Healthy Brain Network (HBN) (16), the Philadelphia Neurodevelopmental Cohort (PNC) (17), the Brazilian High Risk Cohort (BHRC) (18), the Nathan Kline Institute Rockland Sample (NKI) (19), and the Chinese Color Nest Project (CCNP) (20). We used a subset of participants with structural and functional imaging data (n = 3,847; ages 6–22 years; 55.4% male) for the analyses presented. In this dataset, each participant was demonstrated by combination of numerical features derived from brain imaging measurement. The dataset contained structural and functional MRI released at the RBC complete artifact QC threshold – structural scans rated Pass or Artifact and functional scans rated Pass, processed via FreeSurfer v6.0.1, sMRIPrep v0.7.1, and C-PAC v1.8.5.dev1 (2). Analyses in the present study were conducted on the initial release of the RBC data. RBC has since issued an updated release that addresses processing bugs identified after our analyses were completed; we retained the earlier release for consistency with our fitted models and analyses.

The dataset included two outcome-related variables: chronological age and p-factor, a general dimension of psychopathology derived from behavioral assessments. Chronological age served as the primary prediction target, given the well-established relationship between brain-derived features and age. The p-factor variable was retained for downstream analyses to examine whether individuals whose predicted brain age intervals were inconsistent with their chronological age showed differences in psychopathology.

### Image processing

Full details of MRI acquisition parameters, preprocessing and quality control criteria across five cohorts are described in RBC method paper (2). Neuroimaging features were obtained from RBC processing pipeline, utilizing FreeSurfer for structural imaging processing and configurable pipeline for the analysis of connectomes (C-PAC) for functional image processing. All features were segmented using Schaefer-400 atlas, which divides the cortex into 400 regions, then arranged into 7 resting state networks according to the Yeo 7-network atlas (2).

All neuroimaging data passed the harmonized quality assurance criteria implemented by RBC (2), which include structural QC based on expert manual T1 artifact rating and the automated FreeSurfer-derived Euler number, and functional QC based on in-scanner motion (median framewise displacement). We used the RBC complete-artifact release (structural scans rated Pass or Artifact and functional scans rated Pass). Not all participants had both modalities: structural data (cortical thickness [CT] and gray matter volume [GV], 800 features) were available for 4,404 participants, and functional connectivity data (28 features) for 3,847 participants. Retaining only participants with both modalities complete yielded an analytic sample of 3847 participants and 828 features used as input to the machine learning models.

**Table 1.** Sample Characteristics and Neuroimaging Features of the Analytic Cohort.

| Characteristics |  |
| --- | --- |
| **Total participants (n)** | 3847 |
| **Age range** | 6-22 years |
| **Cohort included** | HBN, PNC, BHRC, NKI, CCNP |
| **FC features (Functional Connectivity)** | 28 |
| **Cortical thickness features** | 400 |
| **Gray matter volume features** | 400 |
| **Total brain features** | 828 |
| **BMI, mean (SD)** | 21.01 (5.26) |
| **P-factor, mean (SD)** | -0.09 (1.01) |
| **Male, n (%)** | 2,131 (55.4%) |
| **Female, n (%)** | 1,716 (44.6%) |
| ***Ethnicity, n (%)*** |  |
| **Not Hispanic or Latino** | 2,679 (69.6%) |
| **Hispanic or Latino** | 994 (25.8%) |
| **Missing** | 174 (4.5%) |

| *Handedness, n (%)* |  |
| --- | --- |
| **Right** | 3,295 (85.7%) |
| **Left** | 445 (11.6%) |
| **Ambidextrous** | 62 (1.6%) |
| *Parental Education, n (%)* |  |
| **Complete tertiary** | 1,785 (46.4%) |
| **Complete secondary** | 674 (17.5%) |
| **Complete primary** | 900 ((1).4%) |
| **No/incomplete primary** | 210 (5.5%) |

### Brain age prediction

To construct brain age prediction models, several supervised regression models were compared. Our candidate models included Ridge regression, LASSO, ElasticNet regression, and SVR, all implemented in scikit-learn, and XGBoost quantile regression (21) implemented via the xgboost Python package. Analyses were performed in Python 3.9.13. Neuroimaging features were used as provided in RBC release without applying any additional site or scanner harmonization. These models have been previously used in brain age prediction (5, 12). For each dataset configuration, models were trained using brain derived features as predictors and chronological age as response variable. Model performance was evaluated using root mean square error (RMSE), which quantifies the average magnitude of prediction error in years, and coefficient of determination (R^2^), which reflects the proportion of variance in chronological age explained by the model. All candidate models were compared on each feature set (functional, structural and multimodal); ridge regression was subsequently used as the base model for the primary conformal prediction analyses on the basis of the results reported below figure (2).

### Construction of brain age intervals

While the standard regression models provide point estimates of age, they do not quantify the uncertainty associated with each prediction. Conformal prediction is a model agnostic framework that converts point predictions into intervals with a specified coverage guarantee (e.g., a 90% interval contains the new observation 90% of the time), without distributional assumptions (14). We applied conformal prediction to generate prediction intervals for brain age.

Given a desired confidence level 1 - α, conformal methods produce prediction intervals that contain the true outcome with probability approximately 1 - α. In this study, a 90% coverage level (α=0.1) was used to construct age prediction intervals. For brain age, this corresponds to chronological age being within the BAI 90% of the time. Three conformal prediction approaches were evaluated: (1) Split conformal prediction (15), constructs prediction intervals by separating the training data into a model-fitting set and a calibration set. The calibration set is used to compute residuals that determine the width of the prediction intervals. (2) The Cross-Validation Plus (CV+) conformal prediction method (15) extends this approach by incorporating cross-validation to improve statistical efficiency. Instead of relying on a single calibration split, CV+ aggregates information across multiple folds, resulting in more stable and reliable prediction intervals. (3) Conformalized quantile regression (CQR) (22) combines quantile regression with conformal calibration to directly estimate upper and lower prediction bounds. All conformal prediction methods were implemented using the brain age prediction models to generate individual-level BAIs.

### Conformal Prediction Methods

To evaluate the conformal inference methods, the dataset was divided into training, calibration, and test sets following the procedure illustrated in Figure 1.

**Figure 1.**
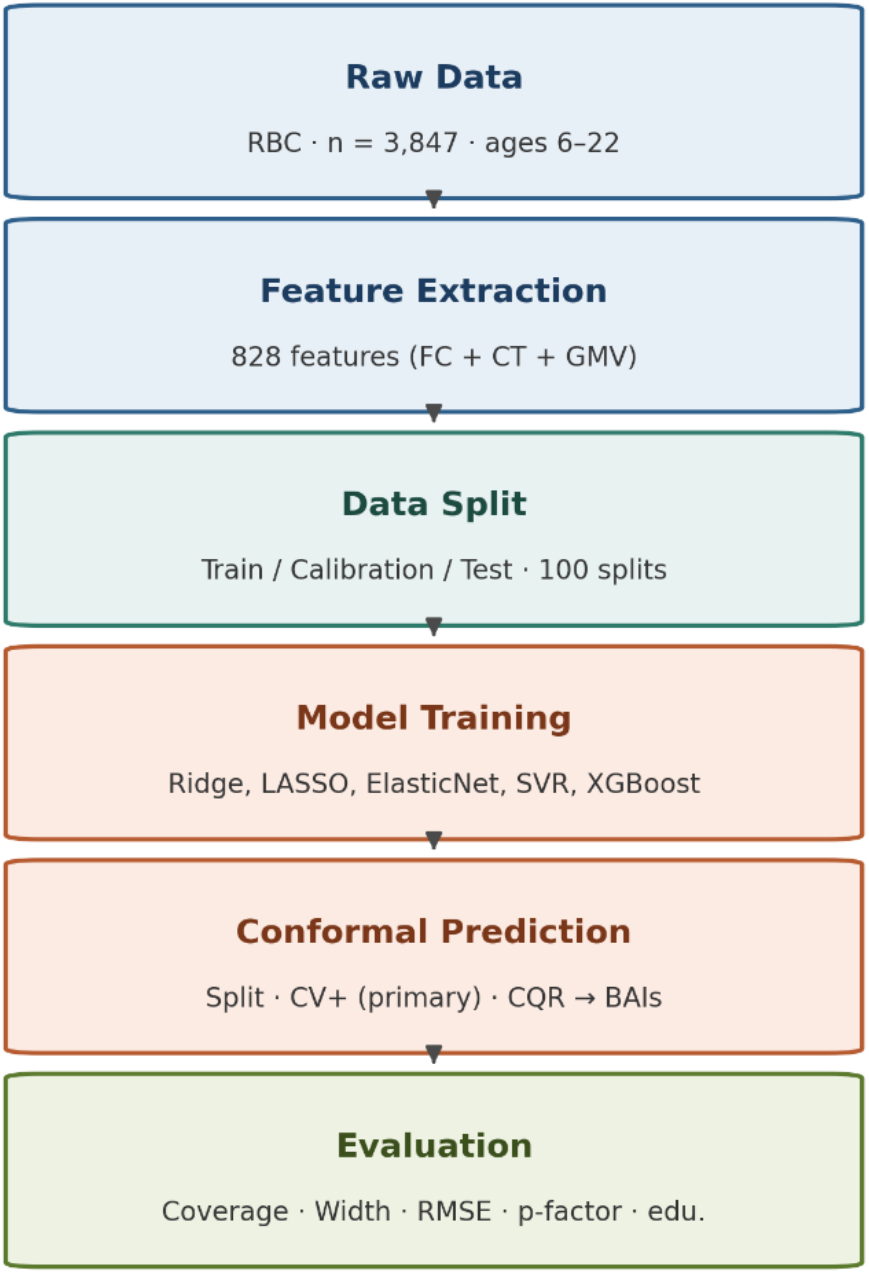
Overview of brain age prediction and conformal inference framework. FC: functional connectivity; CT: cortical thickness; GMV: grey matter volume; BAI: brain age interval; RMSE: root-mean-square error.

For split conformal prediction, the dataset was divided into three sets: 60% for training, 20% for calibration and 20% for test (15, 22). All participants were included in training regardless of psychopathology status; we did not restrict training to a healthy subsample. The training set is used to train regression models while the calibration set is used to determine the prediction interval width. The test data then used to evaluate coverage and interval properties. For CV+ conformal prediction, approximately 80% of the data were used as the training set, which was partitioned into 5 folds. A separate model was trained on each fold by leaving one-fold out at a time. The residuals obtained from the held-out folds were used as nonconformity scores. For each test observation (20% of the data), predictions from all fold models were aggregated to construct the final conformal prediction interval. For CQR, we used the same 60/20/20 split of the data (training/calibration/test) as in split conformal prediction. The training set was used to estimate conditional quantiles of the response variable, and the calibration set was used to adjust the quantile estimates to ensure valid coverage. The resulting prediction intervals were then evaluated on the test set.

### Validation across random splits

Entire analysis was repeated across multiple random data splits to assess the robustness and stability of the conformal prediction framework. For each repetition, the dataset was randomly partitioned following the same procedure described above, and prediction intervals were constructed using the CV+ conformal prediction method with Ridge regression as the base model. Key performance metrics including empirical coverage, average interval width, and RMSE were calculated on the test set for each split. Results across all repetitions were summarized using the median and interquartile range (IQR). This repeated-split validation allowed us to evaluate the stability of the conformal prediction intervals and the consistency of the observed relationships between age prediction intervals and p-factor. We further evaluated empirical coverage across age ranges by partitioning the test set at the median chronological age and computing coverage separately for participants with ages below the median and those at or above the median.

### Associations with p-factor and parental education

To evaluate differences in p-factor and parental education attainment across interval groups, we employed three participant groups based on each individual’s position relative to their predicted 90% BAI (defined earlier): covered participants whose chronological age fell within the interval, uncovered participants with ages above the prediction interval (older group), and uncovered participants with ages below the prediction interval (younger group). We first compared covered versus uncovered participants using Kruskal–Wallis. We then performed pairwise comparisons between covered, older, and younger groups using Mann-Whitney U tests. Statistical significance was evaluated using a threshold of p < 0.05. All analyses were performed using Python using scipy.stats. Within each split, p-values across 2 contrasts (old vs covered + younger, young vs covered + older), were adjusted using Benjamini-Hochberg correction because the covered group is shared across both tests. The p-factor analysis used this framework without additional preprocessing; parental education analysis required additional steps below.

We also explored whether parental educational attainment was related to brain age interval coverage and to the brain age gap. Education was measured on a four-point ordinal scale ranging from no or incomplete primary education (0), complete primary (1), complete secondary (2), complete tertiary education (3); We used the maximum across both parents as our parental educational attainment measure. Across 100 repeated train–test splits, we compared median parental education between older brain group and rest of the sample (covered + younger) using Mann–Whitney U tests, with rank bi-serial correlation as the effect size. As a complementary analysis using traditional brain age gap framework, we regressed the brain age gap on parental education in each split while controlling for chronological age using ordinary least square regression and tracked both significance and direction of the education coefficient across splits.

## Results

We first present results for our model comparison and selection (4.1), coverage of the resulting intervals (4.2), then the relationship between interval membership, psychopathology (4.3-4.4), and parental educational attainment (4.5).

### Comparison of Brain Age Models and Feature Sets Across Repeated Data Splits

CV+ conformal prediction consistently provided the best balance between empirical coverage and prediction interval width across datasets and was therefore selected as the primary conformal framework. Among base regression models compared, ridge regression yielded the narrowest intervals with coverage close to the nominal 90% level across all the 3 feature sets (Figure 2A-F) and was selected as the base model for subsequent analyses. We repeated the CV+ conformal prediction procedure across multiple random 100 train–test splits for each dataset configuration, computing prediction accuracy (RMSE), explained variance (R^2^), empirical coverage of the 90% prediction intervals, and average interval width. Performance metrics were summarized using the median and interquartile range across repetitions.

**Figure 2.**
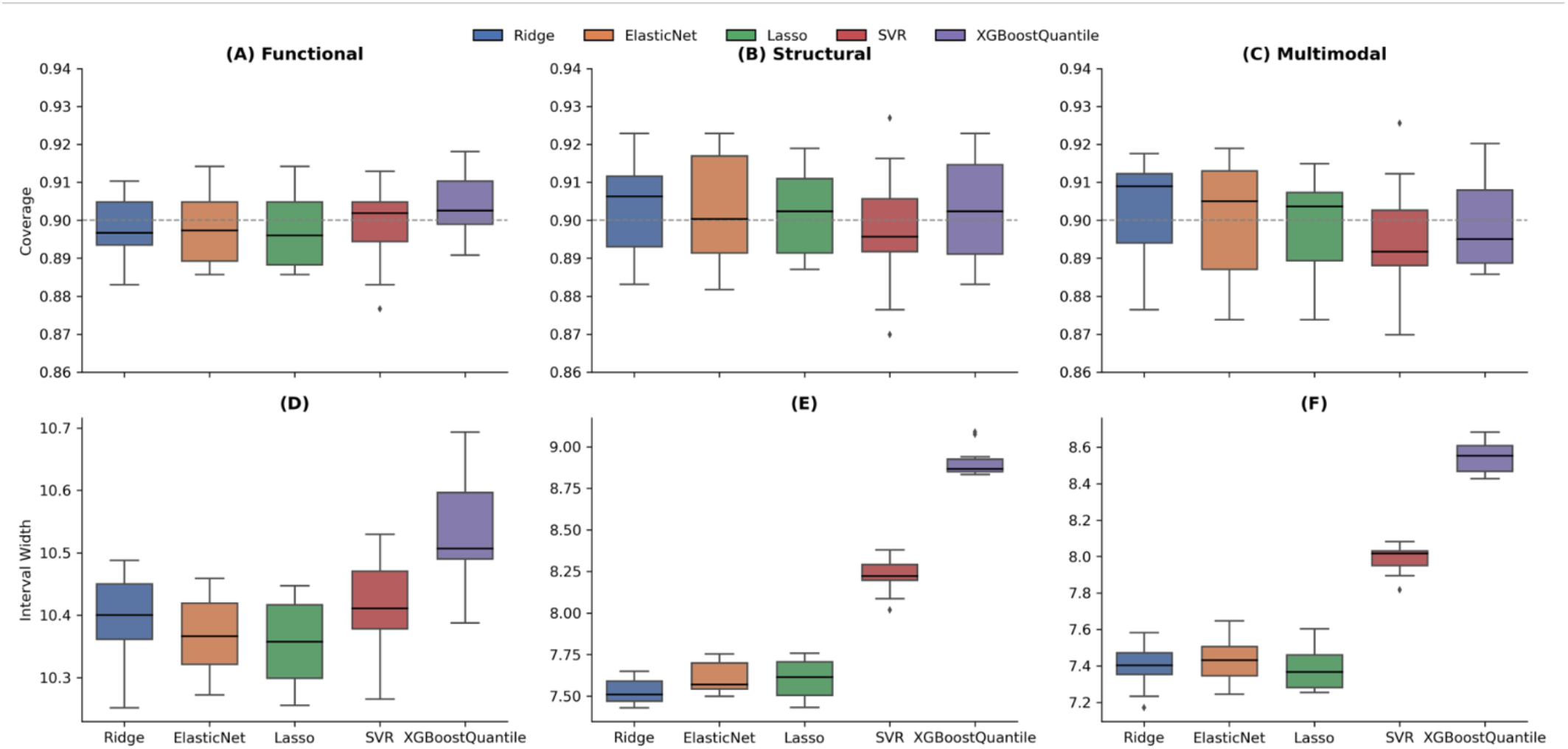
Stability of conformal prediction intervals across feature modalities

Empirical coverage remained close to the nominal 90% target level, indicating reliable calibration of the conformal intervals across models. Median coverage values ranged from approximately 0.89 to 0.91 across models, with only limited variability across repeated data splits in the test set. Figure 2 shows the distribution of empirical coverage (panels A-C) and prediction interval width (panels D-F), across repeated train–test splits. The dashed horizontal line indicates the nominal 90% coverage level.

Throughout all feature sets, structural and multimodal features generated significantly narrow intervals and reduced RMSE in comparison to functional connectivity models. Median interval width for functional models ranges from approximately 10.40-10.81 years for the functional models (Figure 2D), 7.45-9.75 years for structural models (Figure 2E) and 7.30-9.75 years for multimodal models (Figure 2F). CV+ with RidgeCV applied to the multimodal feature set was selected as the primary configuration for subsequent analyses, achieving median empirical coverage of 0.907 (IQR 0.899-0.916) and median interval width 7.36 years (IQR 7.31-7.41).

### Coverage across age groups

One of the key questions we wanted to address is whether the conformal prediction intervals perform equally well across all ages, or whether coverage holds up better for some developmental stages than others. While conformal prediction guarantees coverage at the level of the full dataset, it makes no promises about specific subgroups — so we examined this empirically by computing coverage within two-year age bins across all 100 repeated train–test splits (Figure 3A).

**Figure 3A.**
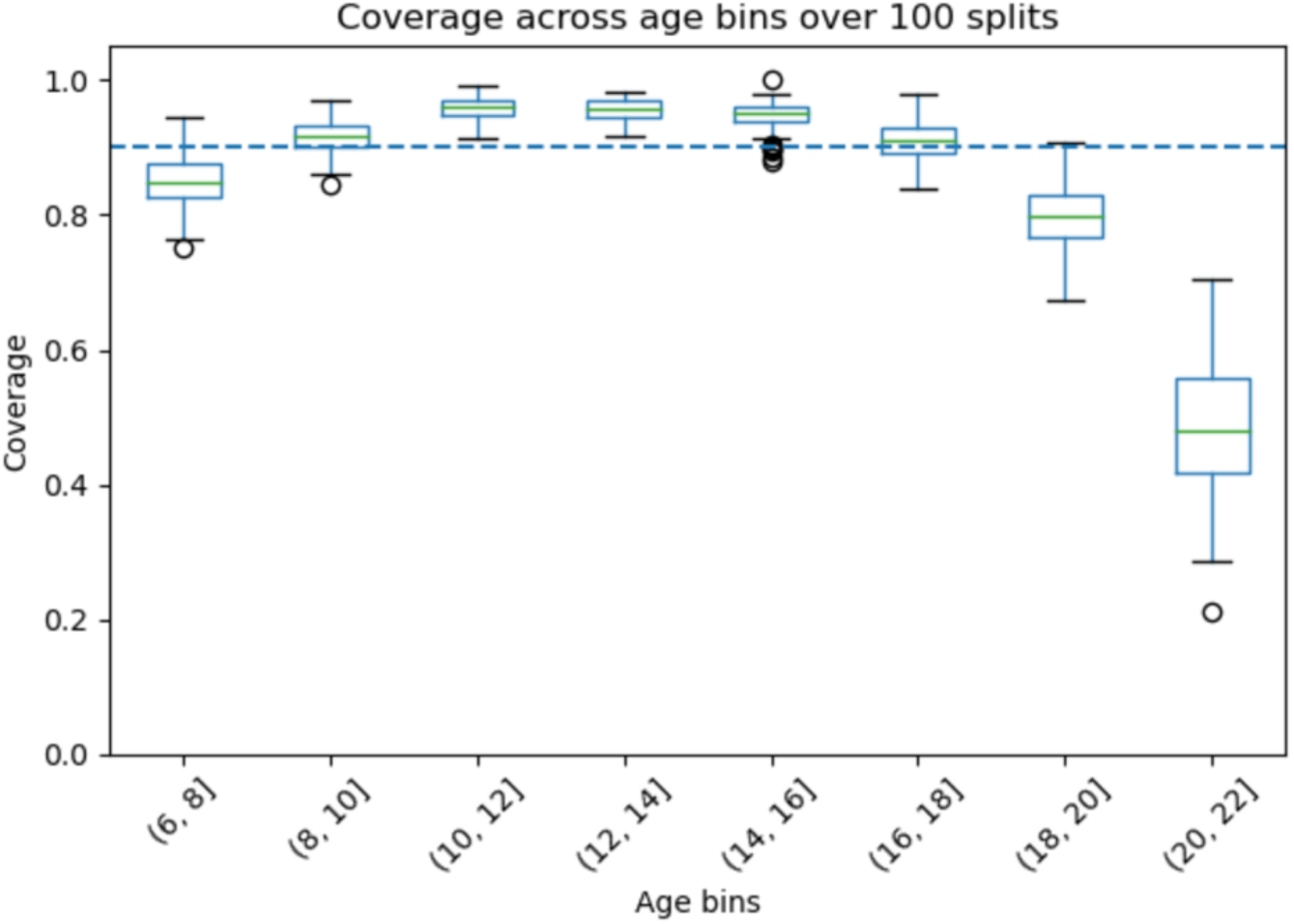
Empirical coverage of CV+ prediction intervals across two-year age bins, computed over 100 repeated train–test splits. The dashed line indicates the nominal 90% coverage level.

**Figure 3B:**
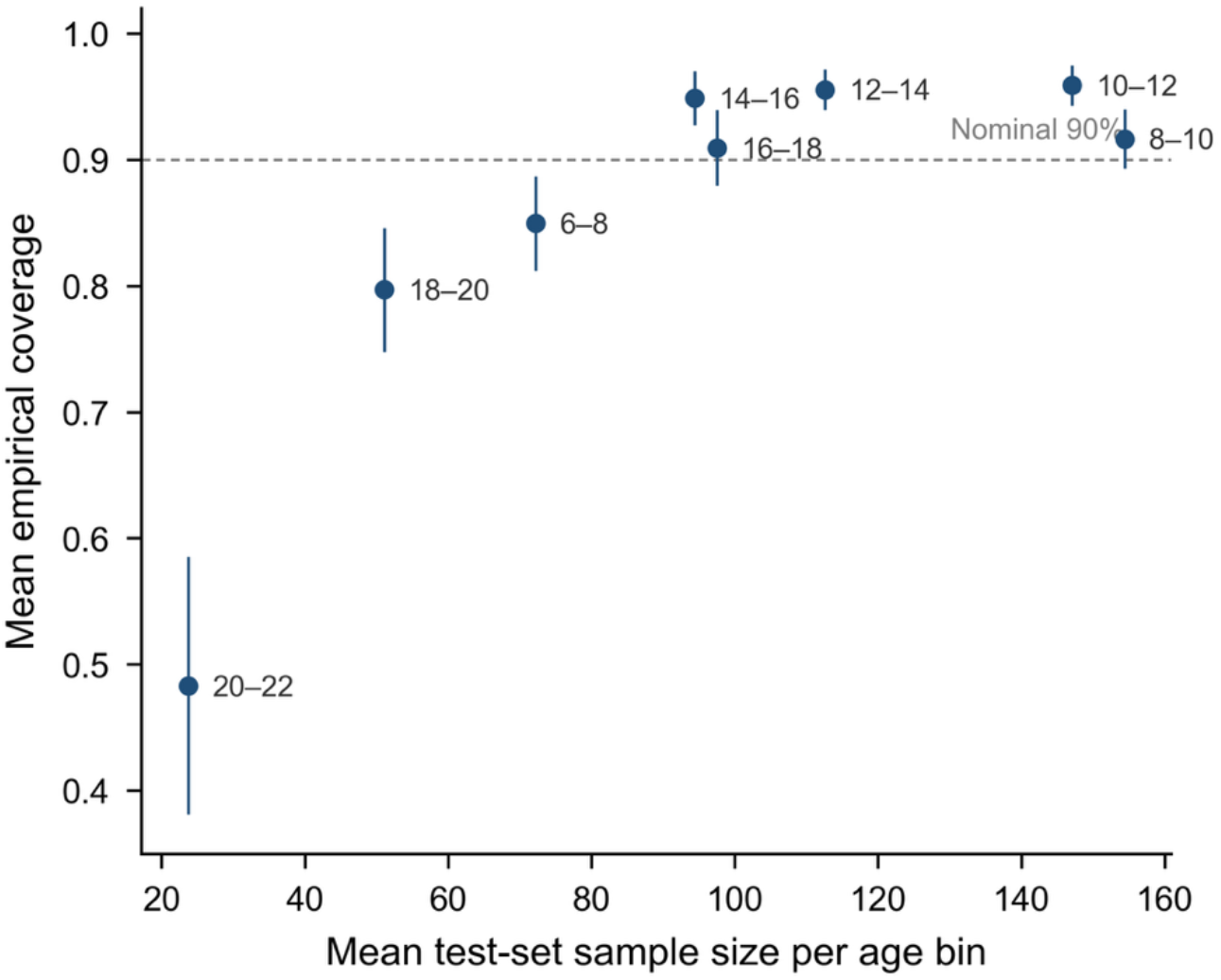
Coverage versus sample size across age bins for a representative split.

**Figure 3C:**
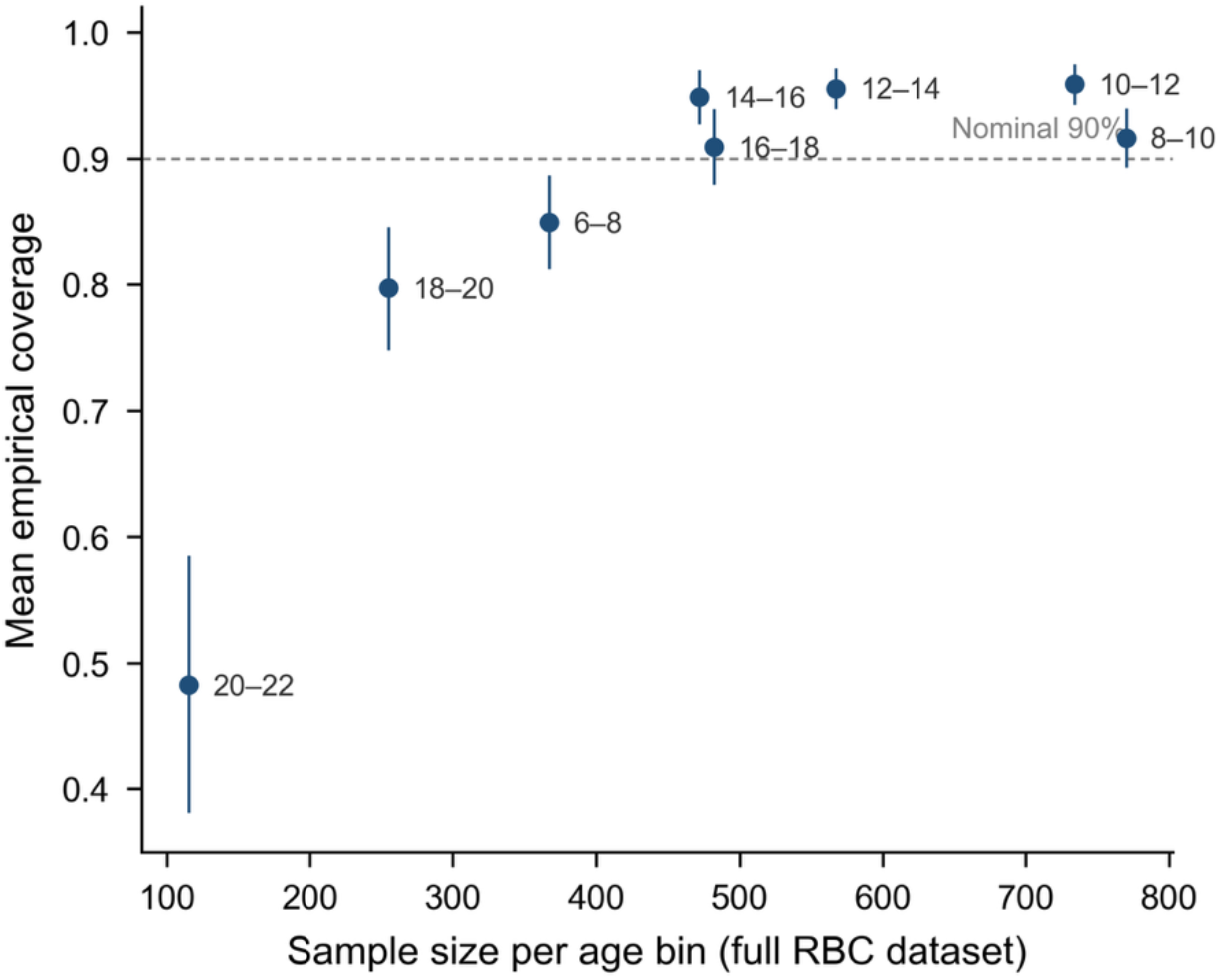
Coverage versus sample size across age bins for a full RBC dataset.

Among youth, Coverage was highest and most stable in the 10–12 and 12–14-year bins, where mean coverage across splits reached 95.9% and 95.6% respectively, both comfortably above the 90% nominal target. The 14–16 and 8–10-year bins followed closely at 94.9% and 91.7%. Even the 16–18 bin held close to the nominal level at 90.9%. So, for the bulk of the sample — participants roughly between ages 8 and 18 — the intervals were well-calibrated and reliable.

In adults, coverage dropped to 79.7% for individuals aged 18–20 and fell sharply to just 48.3% for those in the 20–22 range. This is substantially below the intended level, and the spread across splits was also much wider in these bins, meaning the coverage was not just lower on average but also less predictable from one split to the next.

Looking at a representative split, only 44 test participants fell in the 18–20 bin and 25 in the 20–22 bin — together accounting for under 10% of the full test set. The training set showed a similar pattern, with just 176 and 100 participants in those same bins out of 3,009 total (roughly 9% of training data). When a model sees so few adults during training and calibration, it has little basis for setting appropriate interval widths for that age range, and the coverage suffers as a result (Figure 3B).

This relationship evident when we plot mean coverage across 100 splits against total sample size per age bin in the full RBC dataset (Figure 3C). Bins with fewer participants – 18-20 and 20-22 groups – show substantially reduced coverage and greater variability across splits, whereas bins with hundreds of participants sit at or above nominal 90% target.

### Absence of p-factor Associations Across Repeated Splits

We next examined whether interval group membership was associated with differences in psychopathology.

The p-factor analyses were repeated across multiple random train–test splits using the CV+ framework to evaluate the stability of the observed findings. This procedure allowed us to assess whether the absence of associations between interval inconsistency and psychopathology was consistent across different data partitions rather than being driven by a single split of the dataset.

Across repeated splits, the conformal prediction intervals maintained stable predictive performance. The median empirical coverage across splits was 0.907 (interquartile range: 0.899–0.916), closely matching the nominal 90% target level. Similarly, the median prediction interval width was 7.36 years (IQR: 7.31– 7.41), and the median RMSE was 2.20 years (IQR: 2.15–2.24), indicating consistent model performance across splits. We next examined the stability of p-factor differences across groups. Across repeated splits, the median difference in p-factor between uncovered and covered individuals remained close to zero (median difference = 0.010). Similarly, comparisons between directional inconsistency groups showed minimal differences, with median p-factor differences of −0.042 for older vs covered individuals and 0.081 for younger vs covered individuals (Figure 4B).

**Figure 4.**
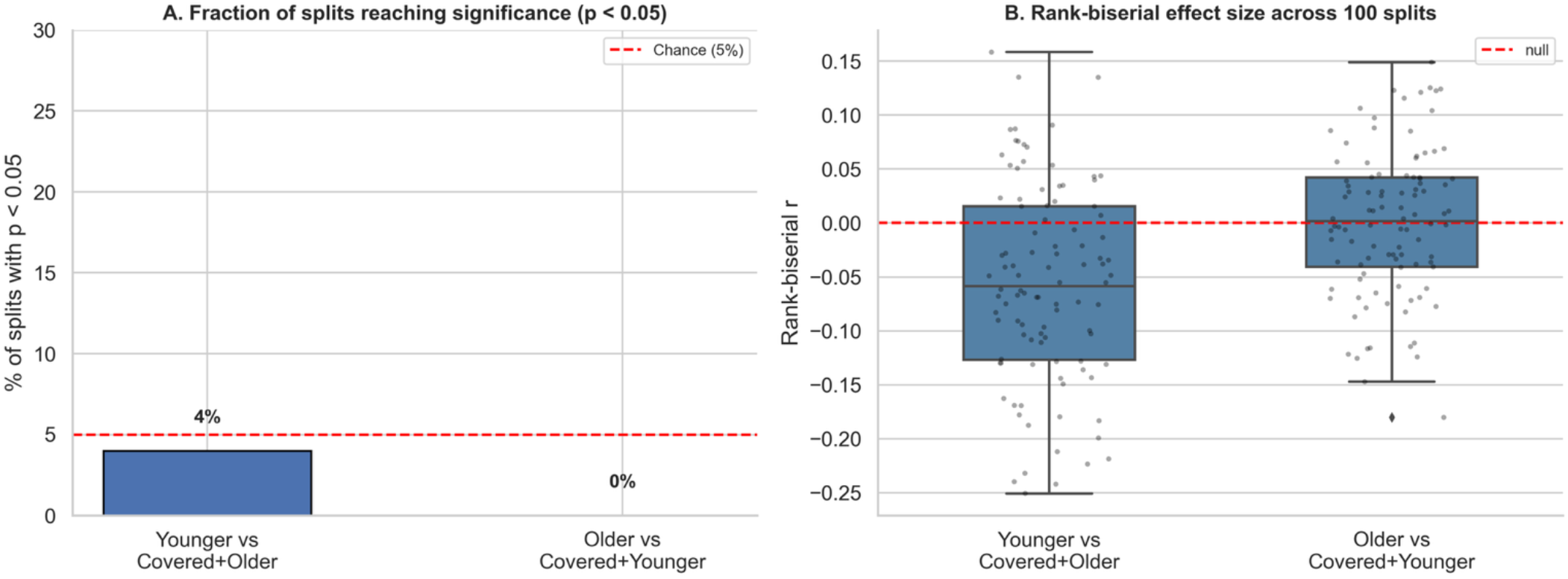
p-factor shows no association with brain age interval coverage across 100 train–test splits.

Statistical tests conducted across repeated splits rarely yielded significant results: the older vs covered + younger Mann Whitney contrast reached p < 0.05 in 0 out of 100 splits, the younger vs covered + older contrast in 4 out of 100 splits (Figure 4A). After Benjamini-Hochberg correction across contrast within each split, 0/100 remained significant for older vs covered + younger and 1/100 for younger vs covered + older, indicating that the absence of an association between interval membership and p-factor is robust to sampling variability. Having established the robustness of the p-factor findings, we next examined whether parental educational attainment showed any association with brain age gap across the same repeated splits.

### Associations between parental education and brain age intervals

Beyond psychopathology, we explored whether parental educational attainment was related to how well the brain age intervals captured an individual’s true age and whether it was associated with brain age gap itself.

Lower parental education was consistently observed in older brain group (older vs covered vs younger) compared to rest of the sample. Across 100 splits, older brain group vs rest contrast produced a weak but directionally consistent effect: median rank-biserial effect size of r = 0.114 (5^th^ - 95^th^ percentile interval [0.002, 0.251]), with lower parental education in the older brain group (Figure 5). The proportion of splits reaching uncorrected p < 0.05 (19 out of 100) is descriptive rather than a formal test, as split resamples of the same participants are not independent. After Benjamini—Hochberg across two contrasts within each split, 9/100 splits remained significant for the older brain contrast and 5/100 for the younger brain contrast. In contrast, younger brain group vs rest was not directionally consistent: median rank-biserial correlation r = 0.031 (5^th^ - 95^th^ percentile [-0.104, 0.204]), with effect size band spanning zero.

**Figure 5.**
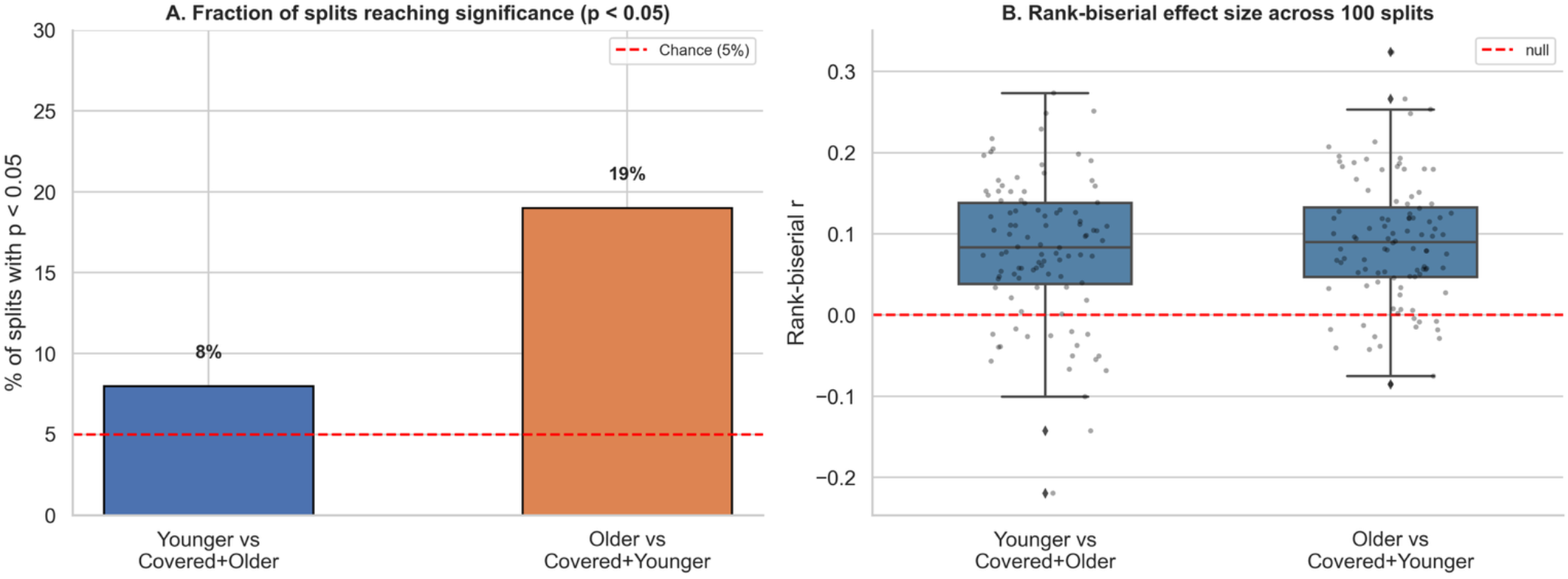
The older brain group shows lower parental education across 100 train–test splits.

For each of the 100 splits, we regressed the brain age gap on parental education while controlling for chronological age, which allowed us to directly compare what the interval-based and the conventional BAG frameworks tell us about socioeconomic associations. Under ordinary least squares regression, the coefficient was negative in nearly every split (average β = −0.088), meaning that children from higher-education backgrounds tended to have smaller brain age gaps. The fact that both the interval-based and the BAG-based analyses point in the same direction but fall short of consistent significance suggests that socioeconomic patterning in brain age, at least as captured by parental education in this dataset, is at best a subtle signal that will likely require larger and more socioeconomically diverse samples to detect reliably.

## Discussion

Prior work has begun to apply conformal prediction to brain age in adult samples (23, 24), demonstrating the feasibility of uncertainty quantified intervals predictions on structural MRI features. The present study extends this direction in three respects: it applies conformal prediction to a large developmental sample spanning childhood to early adulthood, compares multiple base regression models and conformal frameworks on multimodal (structural and functional) features and uses interval membership itself - rather than brain age gap – as the unit of downstream comparison with phenotypic measures. In this work, we proposed an uncertainty-aware framework based on conformal prediction to quantify the reliability of individual brain age predictions using neuroimaging features. By applying conformal prediction to machine learning models trained on functional and structural neuroimaging data, we constructed brain age intervals (BAIs) that provide statistically valid coverage guarantees for individual-level predictions. The conformal prediction framework produced prediction intervals that attained empirical coverage near the nominal level across repeated data splits, indicating stable and well-calibrated uncertainty estimates. Interval widths remained narrow and consistent, and coverage was well-maintained across biological sexes and choice of brain age model for children and adolescents. We further examined whose chronological age fell outside the predicted interval showed differences in psychopathology and parental education. Associations were weak overall. Parental education showed a modest directional signal, with those older than the interval having parents with shorter education, and we did not find an association with the p-factor. In summary, our findings demonstrate that uncertainty-aware brain age modeling via conformal prediction offers a reliable and interpretable complement to traditional point-estimate approaches, with some sensitivity to demographic factors, warranting further investigation in youth populations.

Prior brain age work has largely emphasized point estimates and interpreted the brain age gap (BAG) as a summary measure of deviation from normative age (3, 25, 26). Point estimates, however, provide no measure of uncertainty and brain age models have behaved differently across datasets, features sets and modeling choices (10, 12, 13). Our work extends this literature in two ways. First, wrapping standard brian age regression models in a conformal prediction framework produced valid individual level prediction intervals with near-nominal empirical coverage in a large youth sample, without requiring modification of the underlying models. Second, brain age intervals can be interpreted as a normative interval, offering a framework for relating deviations from normative patterns to phenotypic and demographic variables.

Prior literature has indicated a heightened brain age gap in various mental health conditions, such as major depressive disorder and schizophrenia, suggesting that irregular brain aging patterns may define specific psychiatric groups (3, 7, 26). These findings have led to growing interest in the potential use of brain age metrics as transdiagnostic biomarkers of brain health. However, we found minimal distinctions in psychopathology between individuals whose chronological ages fell inside versus beyond the anticipated brain age ranges. This finding indicates that variations from expected brain age ranges might not directly relate to differences in psychopathology among the examined group. Brain age measures capture broad structural characteristics of brain aging rather than disorder-specific neurobiological mechanisms, and this could be one potential reason. While abnormal brain age patterns have been reported in clinical populations, the strength of these associations may depend on factors such as sample characteristics, disease severity, and imaging features used for model training (25, 27). Consistent with this work, recent work has argued that models trained to predict chronological age may not be the most sensitive approach for detecting psychopathology, since age prediction and psychiatric prediction are related but distinct objectives (14). This null result may partly reflect study design: RBC includes primarily community and help seeking samples with modest symptoms severity and participants with psychopathology were retained in training rather than held out at test cases.

### Limitations and Future Directions

Several limitations of the present study should be acknowledged. Initially for training and assessing the brain age prediction models, the results are contingent upon the traits of the dataset. Model performance may be impacted when applied to independent cohorts due to neuroimaging datasets frequently varying in demographic composition, acquisition methods, and population traits. Tackling this variability continues to be a significant challenge for creating reliable brain age biomarkers. Most brain age modeling techniques pertain to cross-sectional analyses, which is another limitation. While predicted brain age can capture deviations from normative aging patterns, longitudinal data are necessary to determine how individual brain age trajectories evolve over time and how they relate to long-term cognitive and health outcomes (6, 9).

A notable limitation concerns age-conditional coverage. Conformal prediction guarantees marginal coverage across the full test set but not within specific age subgroups. Empirical coverage was substantially lower for adults (age ≥ 18) compared to children and adolescents, likely reflecting adults are severely underrepresented in the Reproducible Brain Charts dataset used for model training and calibration data - an issue is related to class-conditional coverage in conformal prediction (11, 15). The framework is therefore well-suited to youth and future work that incorporates larger adult samples, or age-conditional calibration could extend its reliability across the full lifespan. The 90% coverage level used to define interval groups is one of the several plausible choices; future work could extend the framework to multiple coverage level (e.g. 80%, 90%, 95%) for graded rather than binary group definitions.

Parental educational attainment was the only environmental variable examined and other factors – household income, neighborhood characteristics or early life adversity – may provide more comprehensive picture of environmental influences on brain age uncertainty. The multimodal set was heavily imbalanced (800 structural vs 28 functional), so its interval widths are largely driven by structural contributions. Finally, then median interval width of approximately 7.36 years, while providing valid coverage, may be too wider for individual level clinical utility; local adaptive conformal methods or flexible quantile estimator could yield narrower intervals while maintain coverage guarantee.

## Conclusion

In this study, we introduced uncertainty aware framework for brain age prediction by integrating conformal prediction with machine-learning models developed using functional and structural neuroimaging features. These prediction intervals can be interpreted as normative intervals – the range of ages consistent with an individual’s brain features, providing a principled basis for asking whether a given individual falls within expectation. While ensuring consistent predictive performance, our results suggest that adding uncertainty estimation enhances the interpretability of brain age markers. These findings emphasize the promise of uncertainty-aware modeling frameworks to enhance the dependability of neuroimaging-derived biomarkers in upcoming research on brain aging and mental health. Although, conformal prediction offers a statistically sound method for producing prediction intervals (15, 22), additional research is necessary to assess the performance of uncertainty-aware techniques across various neuroimaging datasets and modeling workflows. Future studies should investigate whether incorporating prediction uncertainty improves the identification of individuals at risk and whether such techniques can improve the clinical understanding of brain age biomarkers. Natural next steps include extending the framework to clinical populations and longitudinal designs, which would allow direct tests of whether interval membership at baseline carries prognostic information for later cognitive or psychiatric outcomes. If such associations emerge, interval-based flagging could offer a principled route to translating brain age from group level research metric into an individual level clinical tool.

## Acknowledgements

We thank then participants and investigators of the Healthy Brain Network, Philadelphia Neurodevelopment Cohort, Brazilian High-Risk Cohort, Nathan Kline Institute Rockland Sample and Chinese Color Nest Project for making their data available and Reproducible Brain Charts team for aggregating and harmonizes these resources.

## Data Availability

Data used in this study are publicly available from the Reproducible Brain Charts project (https://reprobrainchart.github.io/) without a data use agreement.

